# Tourism-driven ocean science for sustainable use: A case study of sharks in Fiji

**DOI:** 10.1101/2020.02.04.932236

**Authors:** C A Ward-Paige, J Brunnschweiler, H Sykes

## Abstract

The oceans are in a state of rapid change – both negatively, due climate destabilization and misuse, and positively, due to strengthening of policies for sustainable use combined with momentum to grow the blue economy. Globally, more than 121 million people enjoy nature-based marine tourism — e.g., recreational fishing, diving, whale watching — making it one of the largest marine sectors. This industry is increasingly threatened by ocean degradation and management has not kept pace to ensure long-term sustainability. In response, individuals within the industry are taking it upon themselves to monitor the oceans and provide the data needed to assist management decisions. Fiji is one such place where the dive tourism industry is motivated to monitor the oceans (e.g., track sharks). In 2012, 39 dive operators in collaboration with eOceans commenced the Great Fiji Shark Count (GFSC) to document sharks (and other species) on 592 dive sites. Here, using 146,304 shark observations from 30,668 dives we document spatial patterns of 11 shark species. High variability demonstrates the value of longitudinal data that include absences for describing mobile megafauna and the capacity of stakeholders to document the oceans. Our results may be used to guide future scientific questions, provide a baseline for future assessments, or to evaluate conservation needs. It also shows the value of scientists collaborating with stakeholders to address questions that are most important to the local community so that they have the information needed to make science-based decisions.

## 1. Introduction

The oceans are in a state of rapid change due to climate destabilization and acidification (e.g., see marine heat waves in Oliver et al., 2018), the impacts of which are compounded by the addition of plastics, noise, and other pollutants into an ocean that has already suffered decades of large-scale misuse (Halpern et al., 2015, 2008). In response, some international goals have been initiated to better understand and protect the oceans and the people that depend on them. For example, the Census of Marine Life (COML; 2001-2010) sampled in 80 nations and discovered 6,000 new marine species (www.coml.org); the International Plan of Action for Conservation and Management of Sharks (IPOA-Sharks) aims to ensure that shark catches from directed and non-directed fisheries are sustainable (www.fao.org/ipoa-sharks); the Convention of Biological Diversity (CBD) Aichi Targets aims to protect and restore global biodiversity (https://www.cbd.int/sp/targets/); and the United Nations Sustainable Development Goals (UN-SDGs) aims to conserve and sustainably use the oceans, seas, and marine resources for sustainable development -- the latter two commit to protecting at least 10% of the world’s oceans by 2020 (Aichi target 11 and SDG 14.5). As well, 2021 marks the beginning of the UN Decade of Ocean Science for Sustainable Development, which aims to generate the global ocean science needed to support the sustainable development of our shared ocean (www.oceandecade.org).

The science needed to evaluate or improve these initiatives pose significant challenges. Importantly, these programs tend to be costly and are typically ‘by invite only’. For example, the COML cost $650 million and involved 2,700 scientists, or <0.1% of the 2.7 million marine scientists (www.datausa.io). The role of stakeholders in these programs, as leaders or as citizen scientists, is largely absent. However, this is slowly changing as the importance of stakeholder involvement and value of citizen science generated data has come to light (Hind-Ozan et al., 2017; Ward-Paige et al., 2013; Ward-Paige, Mora, et al., 2010). Now, scientists are advocating for the inclusion of citizen science and citizen scientists in marine science and conservation (Cigliano & Ballard, 2017), including in meeting SDGs (Fritz et al., 2019).

Under the broad umbrella of citizen science, marine tourism is considered an important partner. Individuals in the marine tourism sector regularly visit various coastal and marine ecosystems and encounter many species or threats. A few programs have filled important data gaps and have been found to provide opportunities to promote trust, education, outreach, awareness, and best practices for ecotourism (Hind-Ozan et al., 2017; Lawrence et al., 2016; Ward-Paige et al., 2014). Additionally, marine tourism operators, guides, and tourists are proving to also be highly motivated to document their oceans (e.g., species and anthropogenic threats) and leverage the economic value of their industry towards improved science, management, and conservation for the oceans and their livelihoods (Ward-Paige et al., 2018, and the current study).

Now that nature-based coastal and marine tourism has significant social and economic value, there is an economic rationale to manage the oceans for reasons beyond commercial fishing. Globally, more than 121 million people take part in nature-based ocean activities, such as scuba diving, snorkeling, recreational fishing, and wildlife watching (Spalding et al., 2016). This sector generates more than $400 billion dollars per year (Spalding et al., 2017), rivaling commercial fisheries, aquaculture, or oil and gas in some areas (National Oceanic and Atmospheric Administration (NOAA), Office for Coastal Management, 2019). The value varies by region and market segmentation (Beaver & Keily, 2015), and ecosystem (e.g., coral reefs at $37.8 billion (Spalding et al., 2016), but combine into one of the largest and most valued industries (Spalding et al., 2017; Spalding et al., 2016). Specific species also drive these industries. For example, shark, ray, and turtle tourism attracts millions of people (e.g., scuba divers), generating direct revenues for local operators and businesses, and contributing to economies on regional and nationwide scales (Huveneers et al., 2017; O’Malley et al., 2013; Troeng & Drews, 2004).

Nature-based marine tourism, however, is vulnerable to both acute and chronic anthropogenic impacts on ocean ecosystems, including those driven by tourism itself. Destruction of corals, mangroves, and seagrass meadows by mining, deforestation, pollution, disease, and aquaculture (Carpenter et al., 2008; Polidoro et al., 2010; Waycott et al., 2009; Zaneveld et al., 2016) reduces the potential area that can be explored for tourism, and diminishes essential habitats for many tourism-targeted species. Overexploitation by fisheries threatens animal populations, including those sought by tourists (Dulvy et al., 2014; Lewison et al., 2014; Worm et al., 2013). Climate change associated impacts, such as acidification, rising sea temperatures, and changing physical oceanographic conditions and nutrient cycling reduce diversity and disrupt the spatial and temporal distribution of animals (Hoegh-Guldberg et al., 2007; Pecl et al., 2017), thus decreasing encounters and making them more unpredictable.

High resolution scientific monitoring is resource intensive, and rarely evaluates the areas that are valued by tourism. Government assessments tend to focus on maximizing commercially exploitative industries (e.g., identifying maximum sustainable yield for commercial fisheries), where only broad trends warrant peer-reviewed publication (e.g., Costello et al., 2016). Academically driven marine-ecology work, especially on marine megafauna, often takes place where animals are relatively abundant in remote and underexplored areas (e.g., Block et al., 2011; Bradley et al., 2017). These two traditional monitoring directives, therefore, rarely overlap in space, time, populations, or interest with nature-based tourism industries.

Where there is overlap between research objectives and tourism, some researchers have begun to solicit the help of citizens, which has grown exponentially in recent years (Follett & Strezov, 2015). In coastal and marine environments, citizen science has been promoted as an important part of governance and management (Ward-Paige et al., 2014) and sustainable tourism (Lawrence et al., 2016), and a number of projects have been launched with the help of citizens to collect baseline data on the relative abundance and diversity of marine life (Brooks et al. 2011; White et al. 2013; Goetze et al. 2018). Many scientific insights have been gained because of these collaborations. For example, citizen contributed data have been used to document species range extensions due to climate change (Last et al., 2011), exotic species invasions (Côté et al., 2013), large-scale absence of reef sharks in proximity to humans (Ward-Paige, Mora, et al., 2010), and the spatial extent of marine garbage (Jambeck & Johnsen, 2015; van der Velde et al., 2017).

In Fiji, similar to other areas, the dive tourism industry depends on diverse, abundant, and reliable marine megafauna encounters to satisfy visitors. Shark diving alone attracts 78% of the country’s 63,000 visiting dive tourists, and inputs over USD 42 million annually to its economy (Vianna et al., 2011). Diving is offered in all bioregions across Fiji (Vianna et al., 2011; Wendt et al., 2018). Despite the considerable socioeconomic value of the diving industry in Fiji, a paucity of information on the diversity, occurrence, and relative abundance of sharks remains and the community has voiced particular concern for sharks in the region.

In response, there has been recent momentum to improve management policies. These include support for locally managed marine protected areas (Govan et al., 2008; Jupiter et al., 2014), mitigating climate change threats (Wendt et al., 2018), at the United Nations Ocean Conference (UNOC – 2017) the Wildlife Conservation Society’ Fiji Country Program signed up to voluntary commitments to expand marine protected areas (https://medium.com/wcs-marine-conservation-program/fiji-makes-16-major-commitments-to-the-ocean-c6f8efce02cd), and the Ministry of Fisheries and Department of Environment in Fiji has committed to develop a comprehensive Shark and Ray Conservation Regulation that ensures sustainable population levels of sharks (and rays) in Fijian waters as part of the SDGs (www.oceanconference.un.org/commitments/). But, there remains a lack of data and resources to use science-based decision making.

To address these data gaps and growing conservation interest, the Great Fiji Shark Count (GFSC) was launched in collaboration with eOceans (previously eShark) to establish a countrywide contemporary snapshot of sharks from the dive tourism industry. Here, we use these longitudinal observations to describe the first nation-wide patterns of diversity, occurrence, and relative abundance of this megafauna group. These patterns may provide a backdrop for future research questions, as a contemporary baseline to compare future populations and human use patterns against, and to inform management or policy, including where, when, and what species to prioritize in conservation initiatives (e.g., in the design of marine protected areas, or to evaluate locally managed marine areas). Additionally, large-scale participation in the GFSC provides an opportunity to consider the impact these types of crowdsourced projects could have on society and sustainability through community collaboration, education and outreach to broader public, and multi-purpose directives (socioeconomic, cultural, environmental, ecological values), which promote movement towards meeting Sustainable Development Goals (https://sustainabledevelopment.un.org/, e.g., Goal 14), various Aichi Targets (https://www.cbd.int/sp/targets/), while also advancing efforts towards the UN Decade of Ocean Science for Sustainable Development.

## 2. Methods

From 2012 to 2016, 39 dive operators across Fiji commenced the first nationwide, longitudinal underwater visual census of dive sites for sharks as part of the Great Fiji Shark Count (GFSC) in collaboration with eOceans (eOceans.co; Fig. 1, Table 1). Each April and November, divers from participating dive shops recorded the details of every dive on 592 sites in 25 areas into community logbooks, including those where no sharks were observed. Before each dive, dive guides instructed guests about the marine region, the objectives of the GFSC, and presented a field guide to correctly identifying the sharks and were given the opportunity to record the details of their observations in the GFSC community logbook.

**Figure 1.**
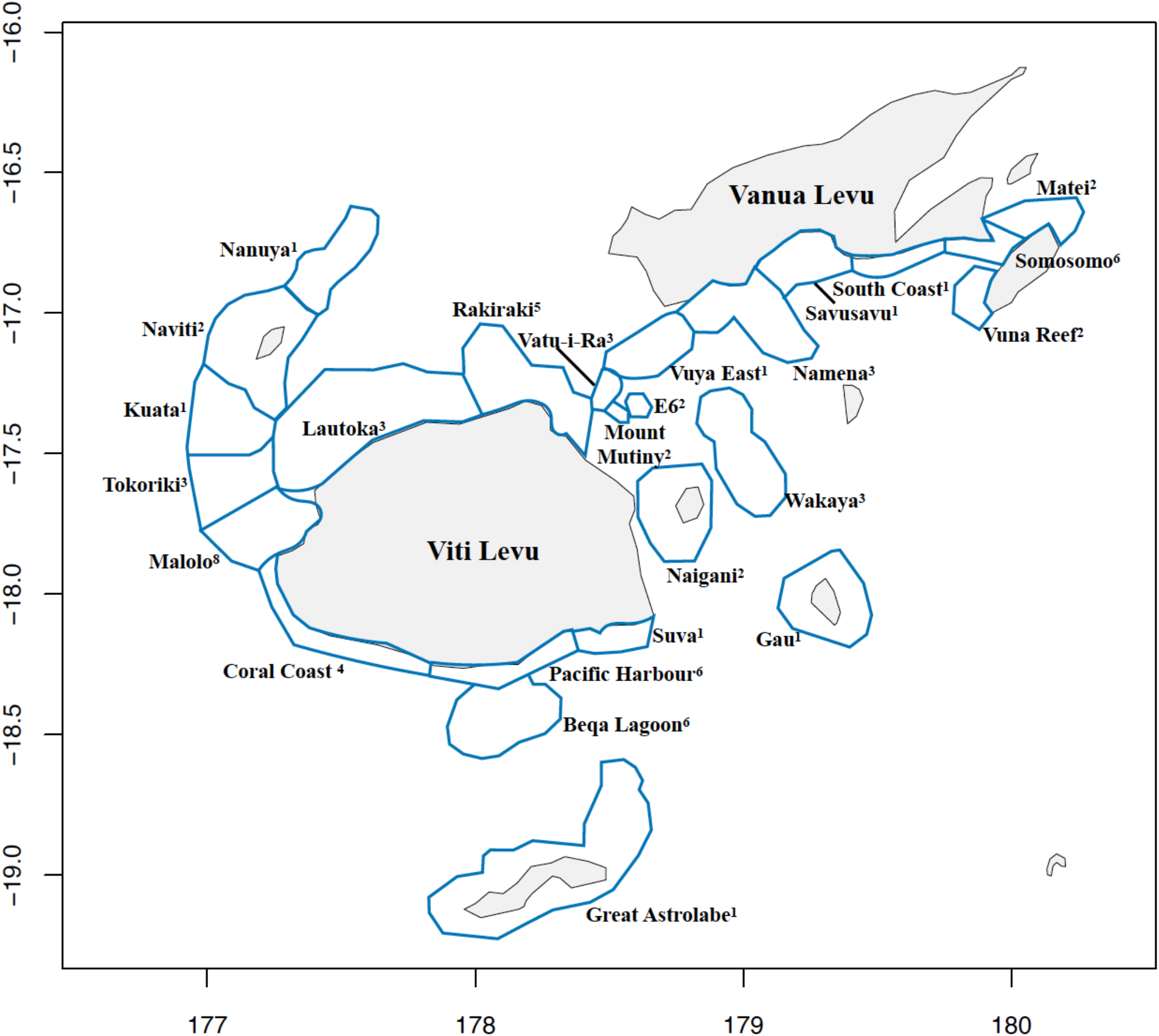
Study region with survey areas sampled by dive tourism operators. Superscripts show the number of operators sampling in each area.

**Table 1.**
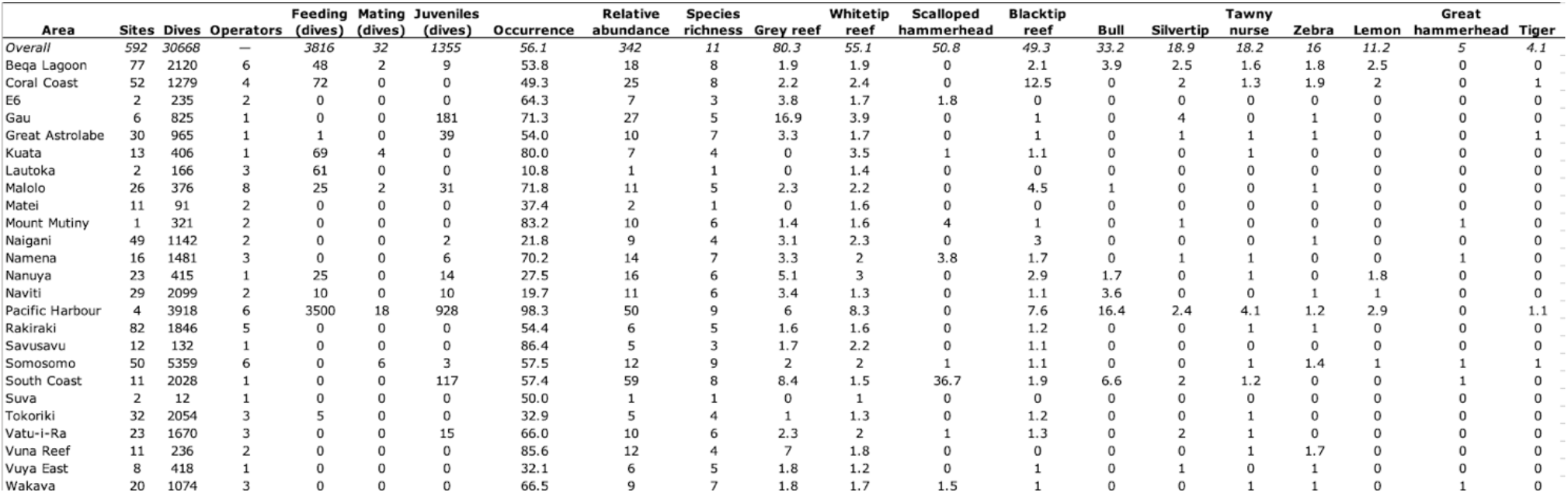
Summary of effort and shark sightings overall and by area. Occurrence and abundance is the mean number without zeros, species richness is the total number of species encountered. For each species in each area, the relative abundance (mean school size without zeros) is shown.

For each dive, including all replicates (i.e., multiple peoples’ observations on the same dive site at the same time), divers logged various attributes of the dive with their observations. Details included: date, time in and time out, operator name, site name, yes or no to spearfishing, and yes or no to wildlife feeding (berleying, chumming, provisioning, etc.). Participants also recorded the presence or absence of sharks (and other species not shown here), and for each species, the number of individuals and if they observed shark mating or numerous juvenile sharks as potential nurseries (Heupel et al., 2007). At the end of each sampled month, all logs were gathered, entered into the national GFSC dataset (fijisharkcount.com), and then into the eOceans database.

To protect sites, species, and communities (e.g., from illegal fishing) sites were assigned to one of 25 areas, without the site coordinates (Fig 1). After the first year, to accommodate differences in site nomenclature between operators, a master site list was created for each area and 207 records were excluded because they could not be matched with a specific site.

For the analyses, we deployed best practices for using recreational divers’ observations at these spatial and temporal scales (Ward-Paige, 2010). We conservatively assumed that all sharks seen by any observer on a site or in area are the same individual sharks seen repeatedly and therefore use the mean number (i.e., average school size) to explore patterns. Operating under these assumptions, three shark metrics were examined: i) richness is the sum of all species observed, ii) occurrence is the percent of dives with sharks or shark species, and iii) relative abundance (abundance hereon) is the mean school size excluding zeros. First, to examine variability at the site-level within and between areas we mapped site-specific effort including number of dives per site and the occurrence or absence of feeds with the three shark metrics. Here, sites were jittered within the area to enable spatial context without compromising site confidentiality. As well, we examined the relationship between effort (i.e., dive count) and the three shark metrics using the loess() smoothing function with ggplot2 in R (Wickham, 2016). Then, we examined the effect of shark feeding and captured a contemporary snapshot of the influence of feeding on sharks by comparing feed and non-feed sites with the three shark metrics and mapped shark occurrence with and without feeding across areas. Next, as a proxy of shark residency, we synthesized effort (i.e., number of dives) with occurrence by species in each area to examine how often a species is observed in an area. Finally, as a proxy of shark behaviour and area use (e.g., schooling, etc.), we synthesized relative mean abundance by species in each area to examine how many individuals may be using each area.

## 3. Results

In total, 30,668 unique dive events were reported from 592 sites in 25 areas in Fiji (Table 1; Fig. 2). Effort was consistent between years (5,139 to 7,018 dives) and months (15,283 in April to 15,385 in November), but varied by site in number of dives (1 to 3,320) and area in number of operators (1 to 8), sites (2 to 82), and dives (12 to 5,359) per area (Table 1; Fig. 1 and 2). Feeding occurred on 15 sites (2.5%) and 3,816 dives (12%). Mating was reported on 32 dives (<1%) on seven sites (1%) and juveniles on 1,355 dives (4%) on 23 sites (4%; Table 1).

**Figure 2.**
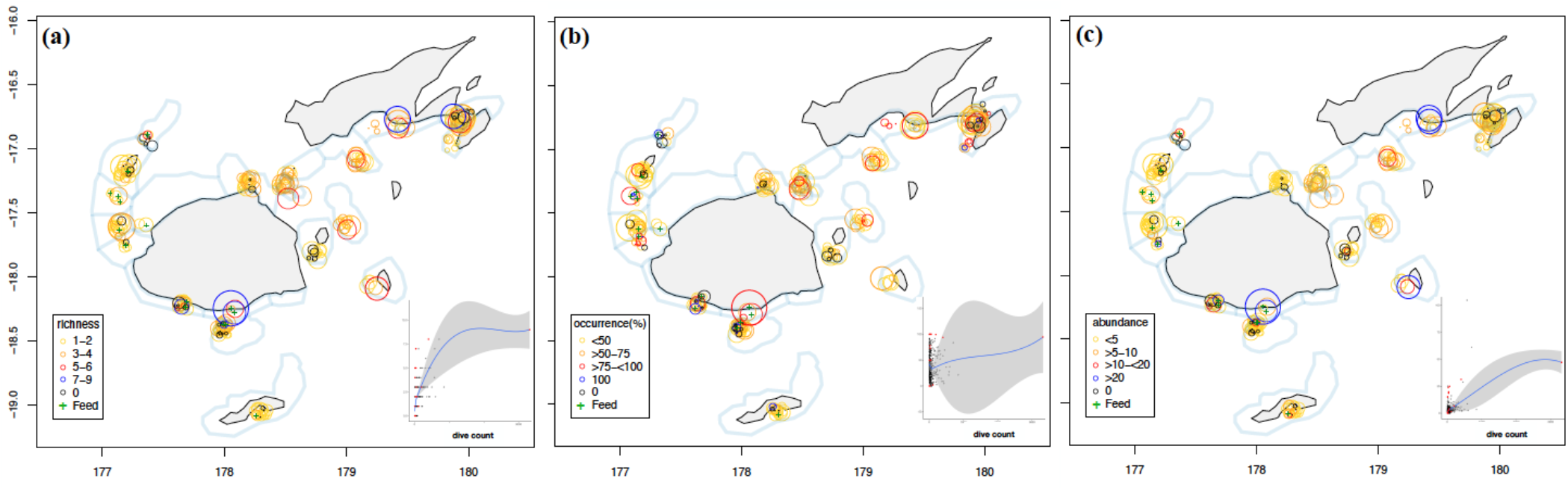
Site specific effort and overall shark occurrence. Shown is shark (a) richness (number of species), (b) occurrence (percent of dives where sharks were encountered), and (c) abundance (mean school size summed across all species). Circle size represents the total number of dives (1-3320) and + are shark feeding sites. For confidentiality reasons, actual site locations are obscured (jittered). Polygons show areas, as specified in Figure 1. Insets show the trend between site-level effort and shark sightings, with feed sites indicated in red and line indicates *loess()* smoothing function.

**Table 2.** Shark species encountered during the Great Fiji Shark Count (GFSC) with Red List Status according to the International Union for Conservation of Nature (IUCN). The number of sharks reported overall, by site, and by area are shown and * show the numbers shown in Figures 2–5.

| Name | GFSC Scientific name | IUCN Red List Status | Total by dive |  | Total by site |  | Total by area |  |
| --- | --- | --- | --- | --- | --- | --- | --- | --- |
|  |  |  | Number | Present | Max | Average* | Max | Average* |
| Whitetip reef | *Triaenodon obesus* | Near Threatened (unknown) | 34,246 | 10,632 | 1,293 | 713.6 | 286 | 55.1 |
| Grey reef | *Carcharhinus amblyrhynchos* | Near Threatened (unknown) | 27,785 | 4,805 | 750 | 334.1 | 359 | 80.3 |
| Bull | *Carcharhinus leucas* | Near Threatened (unknown) | 54,860 | 3,413 | 187 | 55.1 | 136 | 33.2 |
| Blacktip reef | *Carcharhinus melanopterus* | Near Threatened (decreasing) | 13,503 | 2,116 | 320 | 194.1 | 175 | 49.3 |
| Tawny nurse | *Nebrius ferrugineus* | Vulnerable (decreasing) | 3,415 | 935 | 86 | 46 | 61 | 18.2 |
| Scalloped hammerhead | *Sphyrna lewini* | Critically Endangered (decreasing) | 10,626 | 458 | 298 | 166.3 | 161 | 50.8 |
| Lemon | *Negaprion acutidens* | Vulnerable (decreasing) | 890 | 328 | 26 | 14.2 | 23 | 11.2 |
| Silvertip | *Carcharhinus albimarginatus* | Vulnerable (decreasing) | 634 | 286 | 76 | 53.1 | 59 | 18.9 |
| Leopard/Zebra | *Stegostoma fasciatum* | Endangered (decreasing) | 266 | 167 | 37 | 30.1 | 23 | 16 |
| Tiger | *Galeocerdo cuvier* | Near Threatened (decreasing) | 60 | 55 | 13 | 8.7 | 8 | 4.1 |
| Great hammerhead | *Sphyrna mokarran* | Critically Endangered (decreasing) | 19 | 19 | 6 | 6 | 5 | 5 |
| **Total** |  |  | **146,304** | **23,214** | **3,092** | **1,621.30** | **1,296** | **342.1** |

A total of eleven shark species were recorded on 13,846 dives (45%) with a total of 146,304 shark observations made across all dives (Table 2). When sharks were seen, the median and average school size was 3 and 10.6 (SE ± 0.14), respectively, and after accommodating for duplicates within sites and areas and conservatively (using the average school size rather than the maximum) the total number of sharks was 1,621 or 342 individual sharks, respectively (Table 2).

Effort and sightings varied by site within and between areas (Fig. 2). Sharks, of any species, were present on 12,846 dives (45%). Highest effort was on Shark Reef Marine Reserve (Pacific Harbour) with 3,320 dives, followed Tokoriki Wall (Tokoriki), Dream House (South Coast), Nuku Reef (Somosomo Straits), Fish Factory (Somosomo Straits), and Turtle Alley (South Coast) with 572-847 dives. Sharks were present on 441 sites (75%).

Detection of new shark species (i.e., rarer species) on a site generally increased with effort on a site to a maximum of 9 species (Fig. 2a inset). Six sites had seven or more species, including Arena (Beqa Lagoon), Lone Tree (Coral Coast), Fish Factory (Somosomo Straits), Bistro (Pacific Harbour), Dream House (South Coast), Shark Reef Marine Reserve (Pacific Harbour), and Mount Mutiny (Mount Mutiny) had 6 species (Fig. 2a). Shark abundance was highest in Turtle Alley and Dream House, both in South Coast, with 104.7 individuals which consisted of schooling species, including scalloped hammerhead, grey reef, bull, whitetip reef, silvertip, and a few whitetip reef, blacktip reef, and great hammerhead sharks. Bistro and Shark Reef Marine Reserve in Pacific Harbour were the sites with the next most abundant sharks, including schools of bull, grey reef, blacktip reef, nurse, silvertip, whitetip reef, and a few lemon, zebra, and tiger sharks. Shark occurrence was minimally impacted by dive effort where sites with low and high number of dives had varying encounter rates (Fig 2b inset), with an initial negative trend on low effort sites. Occurrence exceeded 90% on 86 sites, and the four sites with the highest effort and occurrence were Shark Reef Marine Reserve (Pacific Harbour -- 3320), Bistro (Pacific Harbour -- 402), Moiya Reef (Kuata -- 195), and The Zoo (Somosomo Straits -- 172; Fig. 2b). Shark abundance, sum of the average school size across all species on a site, generally increased with effort (Fig. 2c inset). Abundance was highest on Turtle Alley (105) and Dream House (61), two non-feed sites in South Coast and the next highest sites were Shark Reef Marine Reserve (47) and Bistro (31), two feed sites in Pacific Harbour (Fig. 2c). Feed sites typically had higher than expected species richness, occurrence, and abundance for a given effort, but it was not always the case (Fig. 2 insets, shown in red).

Effort, the number of feed sites and dives, and shark occurrence varied by area throughout the study region (Fig. 3). Fifteen areas (60%) had no feed sites or dives. Four areas, including Great Astrolabe, Tokoriki, Naviti, and Beqa Lagoon had ≤3% feeding, Coral Coast, Nanuya, and Malolo had 5.5 to 6.5% feeding, and Kuata, Lautoka, and Pacific Harbour had the highest feeding rates at 17%, 37%, and 89%, respectively. Feeding influenced the richness, occurrence, and abundance of sharks observed (Fig. 3a,b). Median shark species richness on non-feed sites as 1 and 3 on feed sites, with outliers occurring on non-feed sites including Dream House (South Coast – 8), Fish Factory (Somosomo Straits – 7), Lone Tree (Coral Coast – 7), and Mount Mutiny (Mount Mutiny – 6). Median shark occurrence was 28 on non-feed sites and 94 on feed sites, with no outliers. Median shark abundance was 1.3 on non-feed sites and 15 on feed sites, with outliers on non-feed sites including Turtle Alley (South Coast – 105), Dream House (South Coast – 61), Nigali Passage (Gau – 29), China Town (Beqa Lagoon – 25), Lone Tree (Coral Coast – 16), Fanny Hill (Coral Coast – 15), Nigali Outside Reef (Gau – 13), Stingray City (Coral Coast – 13), Fish Market (Naigani – 11), Channel (Coral Coast – 11), Soso Passage (Great Astrolabe – 11), SchoolHouse (Namena – 11), Mount Mutiny (Mount Mutiny – 10), Lonetree Channel (Coral Coast – 10), Magic Mushrooms One (Namena – 10), Stairs (Vuna Reef – 10), North Save-a-Tack Passage (Namena – 10), Supermarket (Malolo – 10), Grand Central Station (Namena – 9), Fish Factory (Somosomo Straits – 9), Combe Reef (Pacific Harbour – 8), Frigates (Beqa Lagoon – 8), North Reef (Malolo – 8), and The Zoo (Somosomo Straits – 8). With caution, due to varying sample sizes (15 in feed sites vs. 577 in non-feed), analysis of variance suggests that feed sites are significantly different from non-feed sites for all three metrics (p <0.001). Figure 3b shows shark occurrences mapped by area, with and without feeding. Sharks were most commonly encountered in Pacific Harbour (93%), Savusavu Bay (81%), Kuata (79%), and Vuna Reef (72%), including feed and non-feed dives, and least common in Lautoka and Suva with ≤10% occurrence.

**Figure 3.**
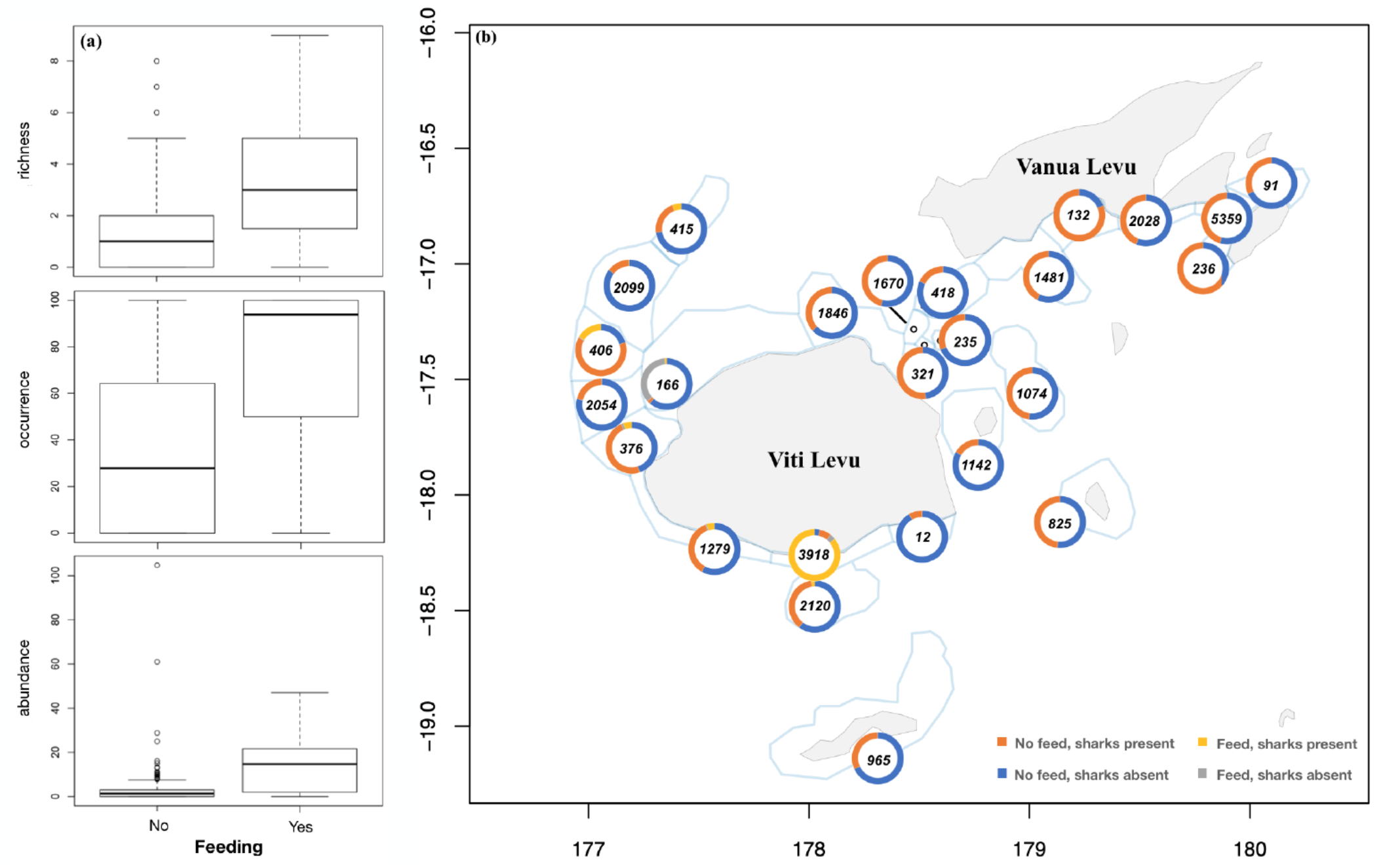
Contemporary snapshot of dive tourism with the prevalence of sharks and shark feeding in Fiji. Numbers show the total effort.

Shark species richness, occurrence, and mean abundance (i.e., school size) varied by area (Fig. 4 and 5). Whitetip reef sharks were the most commonly encountered shark species, being observed in all areas with occurrences ranging from 3% to 80% in Lautoka and Savusavu Bay, respectively (Fig. 4). Grey reef and blacktip reef sharks were also commonly observed. Grey reef were in 21 areas with highest encounters in Pacific Harbour (53%), Gau (44%), and Vatu-i-Ra Lighthouse and Namena (28% each), and blacktip reef sharks in 20 areas highest encounters in Pacific Harbour (37%) followed by Malolo (14%) and Nanuya (13%). Tawny nurse shark was in 14 areas with Pacific Harbour having the highest encounter rate (20%) and the others having ≤3%. Zebra shark was encountered in 13 areas, most commonly in Coral Coast (7%) and Pacific Harbour (5%) with other areas have ≤2%. Silvertip sharks were seen in 10 areas, most commonly in Mount Mutiny (9%) and Pacific Harbour (5%) with other areas having ≤1%. Scalloped hammerhead sharks were in 8 areas, most commonly in South Coast (14%), Wakaya (10%), Mount Mutiny (7%) with all others having ≤3%. Lemon sharks were encountered in 6 areas, most commonly in Pacific Harbour (7%) and Nanuya (6%) with other areas having ≤1%. Bull sharks were also in 6 areas, most commonly in Pacific Harbour (85%) then Nanuya (6%) with others having ≤2%. Great hammerhead and tiger were seen in 5 and 4 areas, respectively, with encounters of ≤1%. Maximum mean school sizes varied by area. The highest mean school size was scalloped hammerhead in South Coast (37). Then, grey reef in Gau (17), Bull in Pacific Harbour (16.4), blacktip reef in Coral Coast (12.5), whitetip reef in Pacific Harbour (8.3), tawny nurse in Pacific Harbour (4), silvertip in Gau (4), lemon in Pacific Harbour (3), zebra in Coral Coast (2), and tiger and great hammerhead sharks with mostly only singles being observed -- the exceptions were two reports of five and two tiger sharks reported in Pacific Harbour on one occasion each (Fig. 5, Table 1).

**Figure 4.**
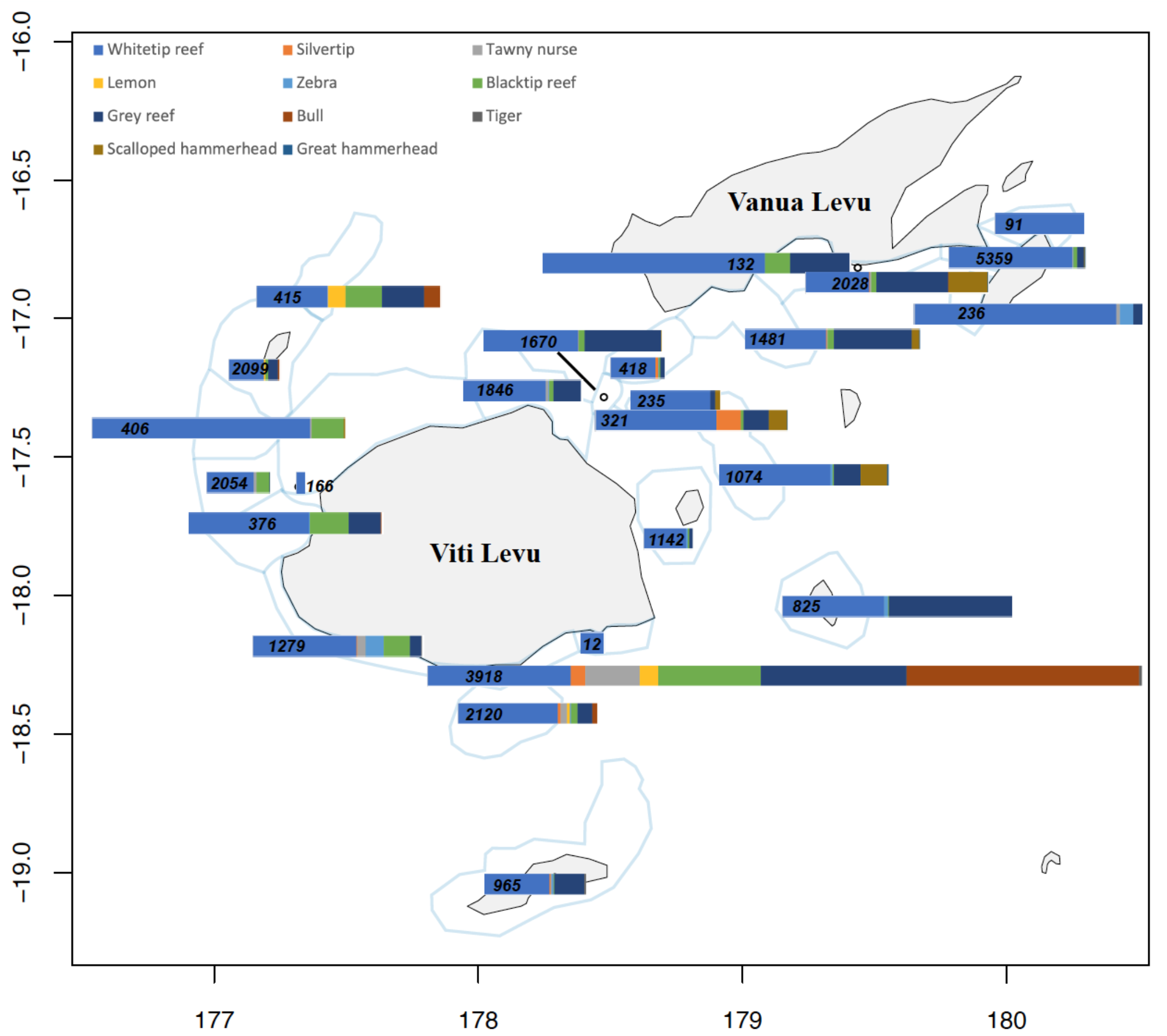
Shark species sighting frequency by area. Segments show the species percentage compared to all shark sightings and segment length is the combined total. Numbers show the total effort.

**Figure 5.**
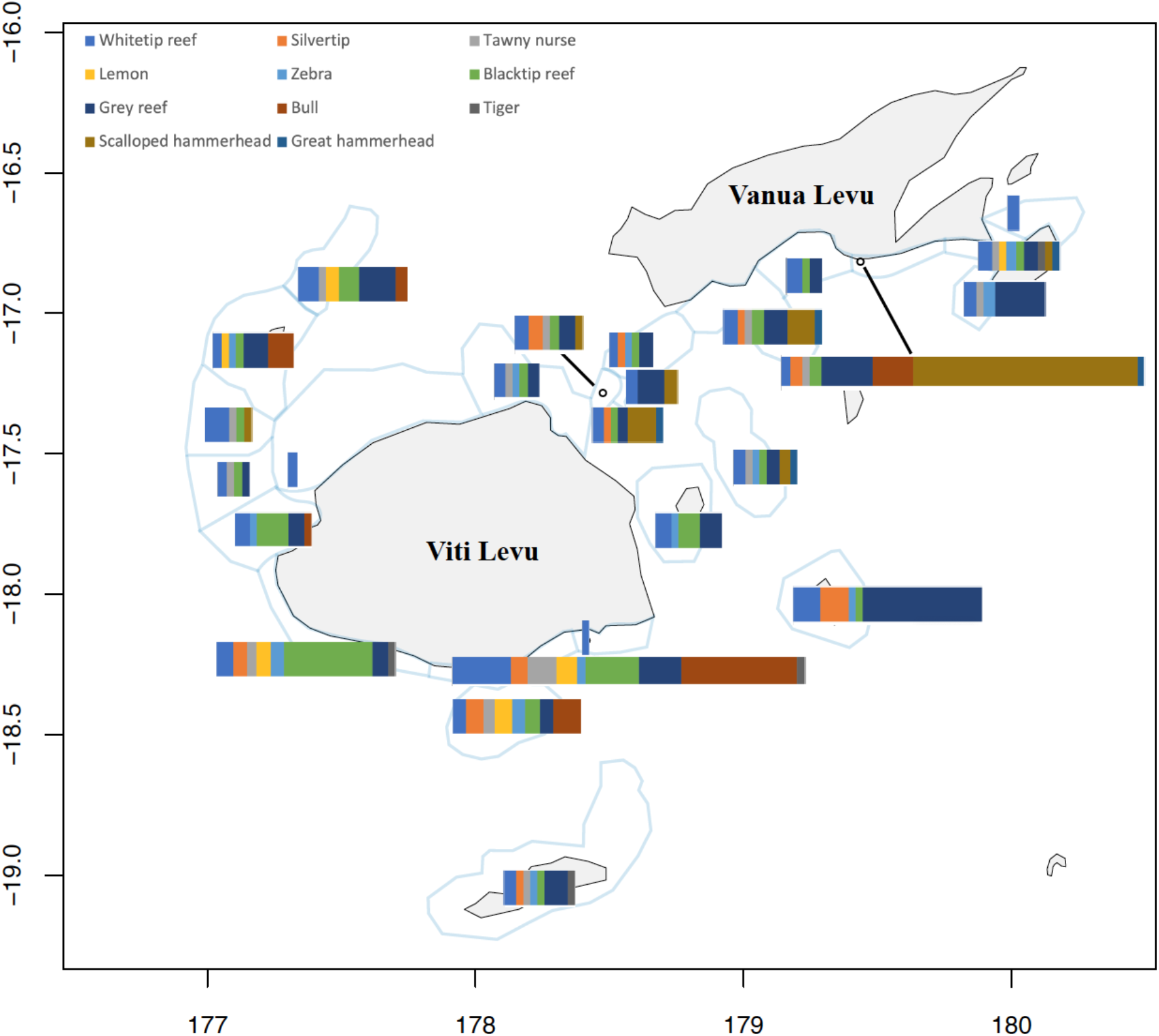
Shark species relative abundance (max/avg school size) by area. Segments show the species percentage compared to all shark sightings and segment length is the combined total. Numbers show the total effort.

## 4. Discussion

### 4.1 Summary

The Great Fiji Shark Count (GFSC) -- a collaboration of 39 dive operators and eOceans -- conducted a nation-wide five-year census of 592 sites across Fiji by collecting observations from >30,000 dives in 25 areas. A total of 146,304 shark observations were used to describe at-sea spatial and temporal patterns of species richness, occurrence, and relative abundance. These species’ distribution patterns in the area are largely undescribed, showing the value of this collaborative program for revealing novel information. All species varied in occurrence and abundance at site and area levels, demonstrating the need for high resolution spatial data that includes observations when no sharks were observed. Feeding elevated shark species richness, occurrence, and abundance, but the effect was site and area specific. Additionally, the long-term, ongoing contribution by dive operators to the GFSC demonstrated high-level of interest by the community to document their activities and ecosystem. Taken together, these observations suggest that the dive tourism industry is motivated to track ocean issues that matter to them, which may serve the broader interests of scientists and decision makers if they find ways to collaborate to improve the speed and accuracy of their own discoveries and to make informed management and policy decisions.

### 4.2 Observed patterns

By reporting every dive, regardless of what was observed (e.g., no sharks), the GFSC was able to gather data to gain novel insights on variations at the species level, which can inform and prioritize scientific research and management.

Our study demonstrated that sharks are seen throughout Fiji and that they are common and abundant enough to be detected by divers in all areas. This is positive since five species are Near Threatened while six species are Threatened -- two Critically Endangered, one Endangered, and three Vulnerable according to the IUCN Red List of Threatened Species (https://www.iucnredlist.org/; Table 2); many have life history characteristics that make them vulnerable to exploitation (Dulvy & Forrest, 2012); many have a long history of overexploitation and are still targeted (S. Oliver et al., 2015; Worm et al., 2013); and, because they are coastal species, are vulnerable to coastal habitat degradation and change (Lotze et al., 2006; Ward-Paige et al., 2015; Waycott et al., 2009). As well, overexploitation and bycatch of sharks has been rampant around the world (Dulvy et al., 2014; S. Oliver et al., 2015; Wallace et al., 2010; Worm et al., 2013), including in Fiji (Glaus et al., 2015), and many are now too rare to be detected by divers in other regions (Ward-Paige, Mora, et al., 2010).

The spatial ecology of the majority of these species has not been described across Fiji before; however, where there is overlap there are mostly minor deviations. Shark species reported in the GFSC are the same as those documented by divers previously (Vianna et al., 2011), and only one species (blacktip shark) was caught by artisanal and subsistence fishers (Glaus et al., 2018) but not reported in the GFSC. Whale sharks were previously documented in northwest Viti Levu on offshore sites (Sykes et al., 2018), but were not recorded in the GFSC. Small numbers of adult scalloped hammerhead sharks have been documented in the Vatu-i-Ra Lighthouse area (Vianna et al., 2011), but our study is the first to record schooling scalloped hammerhead sharks (maximum 12 to 100 individuals) in Mount Mutiny, Wakaya, Namena, and South Coast. A 2004 scientific diver study in the Shark Reef Marine Reserve (SRMR), Pacific Harbour, found that eight shark species use the site (Brunnschweiler & Earle, 2006); with the addition of zebra sharks the GFSC found that the same species still use the site but the mean numbers appear to have changed slightly for a few species with the GFSC having higher blacktip reef (7.9 compared to ~1) and whitetip reef sharks (9.6 compared to ~1), and lower numbers of tawny nurse (1.1 compared to ~4) and grey reef (6.4 compared to ~15) (note: monthly and annual comparisons are needed to be more explicit about change). In Namena, which contains Fiji’s largest no-take marine reserve, a 2009 study using baited remote underwater video systems (stereo-BRUVs) found five shark species (Goetze & Fullwood, 2013); with the exception of zebra sharks (previously a maximum of one individual) the GFSC found that the same species still use the site, with the addition of tawny nurse (max = 1, mean = 1), scalloped hammerhead scalloped hammerhead (max = 30, mean = 3.8), and great hammerhead sharks (max = 1, mean =1). Interestingly, for the species occurring in both studies, the maximum number of individuals seen in the previous study are very similar to the GFSC -- grey reef at 19 and 25 (GFSC mean = 3.3), whitetip reef at 19 and 21 (GFSC mean = 2), blacktip reef at 3 and 3 (GFSC mean = 1.7), and silvertip at 1 and 1, respectively. Further, these similarities between the GFSC diver data and BRUV data, demonstrate the value of high effort dive sampling and how conservative the mean abundance values presented in the current study are.

Tracking and incorporating human use patterns, through the capture of zeros (i.e., where sharks were absent), provides important context for the interpretation of our results. Feeding to attract sharks has become popular in recent decades and has been shown to alter shark behaviour, occurrence, and abundance patterns (Gallagher & Hammerschlag, 2011; Gallagher et al., 2015). Without pre-feed baselines, we cannot fully understand the impact of feeding on shark patterns -- presumably sharks already occurred in higher abundance on these sites compared to other sites but this is not documented. Also, the impact of feeding on sharks in Fiji has been described elsewhere (Brunnschweiler et al., 2014; Brunnschweiler & Baensch, 2011), and is beyond the scope of this study. Regardless, feeding is an important consideration in understanding the contemporary distribution of sharks. Bull sharks, for example, are one of the most sought after species for shark diving tourism in Fiji, (Brunnschweiler et al., 2014) and were reported in six areas during the GFSC. Bull shark occurrence and abundance was highest at two sites in Pacific Harbour, where feeding occurred regularly. Interestingly, however, the area with the second highest abundance of bull sharks was in South Coast (maximum of 20 and mean of 6.6 individuals), which did not report shark feeding. It remains unknown what attracts bull sharks to this area and more importantly if these are the same individuals that can be observed at the SRMR in the Pacific Harbour area (Brunnschweiler & Baensch, 2011). Bull sharks are capable of long-range movements (Brunnschweiler et al., 2010; Heupel et al., 2015) and anecdotal reports show that bull sharks that were visually identified at the SRMR in Pacific Harbour were also recorded in Kuata where a shark feed site was established in 2015 and bull sharks began to be seen late that year (T Vignaud, personal communication) but not reported to the GFSC. Regardless, our results also show that bull sharks are found in variable school sizes, in fed and unfed conditions, further suggesting that feeding has only localized effects on observations. These differences further highlight the importance of tracking and controlling for feeding when describing and monitoring marine animals.

Our results further show the importance of longitudinal sampling for the study of mobile marine fauna, which may have important implications for scientific investigations of mobile marine megafauna more generally. By regularly sampling sites over five years, the GFSC showed clear variations in species occurrence and abundance. Many scientific studies, especially in remote areas, are resource and time limited -- covering a relatively short time period (less than one year) and a small number of sites, often without replicates. For mobile species, even those with relatively small home ranges like reef sharks, these small and short-term censuses may misrepresent populations (e.g., missing species). However, capturing even higher resolution data (e.g., running GFSC all year) could further refine the variation and add additional insights.

Mating and nursery areas are essential habitats (Glaus et al., 2019; Heupel et al., 2007) and are among the important areas to be considered for shark management and conservation (Kinney & Simpfendorfer, 2009). Here, both were only rarely encountered and the locations were variable. Again, conducting even higher resolution censuses, and combining divers’ observations with others’ observations (e.g., artisanal fishers), may help to better identify these areas.

### 4.3 Caveats for observed patterns

There are some general caveats to consider. Sampling was not standardized in space or time, which limits some of the potential analyses and interpretations. As with all visual censuses, visibility, distance to the animal, and diver experience can affect species detection and identification (Darwall & Dulvy, 1996; Thresher & Gunn, 1986; Ward-Paige, Mills Flemming, et al., 2010) and were not addressed. Some species, populations, and individuals may display more or less avoidance behaviour to scuba divers (Cubero-Pardo et al., 2011; MacNeil et al., 2008) including in shark feeding areas (Brunnschweiler et al., 2014), which can impact the observed species and abundance. Because the exact site locations (to protect people and species) were not reported, we cannot estimate the impact of feeding on adjacent sites -- we would expect proximity and species mobility (e.g., mobility) to be an important factor determining the independence of sites (e.g., if the same individuals are observed in nearby or distant sites and areas). When calculating abundance as the mean school size of each species, we assumed that all sightings were repeated sightings. which underestimates the relative abundances, especially for those that are highly mobile or transient and unlikely to be repeatedly observed. Finally, relative abundance is not expected to reflect true abundance, but rather a proxy of the number of individuals observed in each area, and these values need to be carefully considered with human use patterns that may impact shark behaviour (e.g., feeding). Many of these challenges can be overcome by considering population specific information on home range, residency, and site fidelity, which could help to estimate repeat versus independent sightings and refine relative abundance estimates. We do not yet have this level of detail, but tagging studies, such as those done for bull sharks in the Pacific Harbour area (Brunnschweiler & Barnett, 2013), may help to define this further in the future.

### 4.4 Backdrop for future work

This first description of the spatial patterns of these 11 shark species across Fiji lays the foundation for further scientific research, conservation, and management design or evaluation.

Our study provides many novel results that could direct future scientific research. Perhaps with the exception of the bull shark, there is currently not enough published work on these 11 species to be able to interpret many of our findings (e.g., what drives the variation in diversity and abundance). Instead, however, our results provide an initial contemporary baseline that may be used to drive further scientific study. For example, collecting photographs of individual animals could be used to investigate mobility (Araujo et al., 2017; Bradshaw et al., 2007; Couturier et al., 2011; Dudgeon et al., 2008) or threats (Bansemer & Bennett, 2008, 2010) across Fiji.

Further, this five-year snapshot of Fiji’s sharks may be used to evaluate and compare future populations, such as in response to climate change, fishing, or education and management initiatives. In this case, since Fiji has a long history of fishing sharks we cannot presume that our GFSC provides a natural baseline. However, it does provide a contemporary baseline that may be used to compare with future studies. Given the possibility of exploring seasonal or annual trends with GFSC data, and that we have already documented the expected spatial variability and the commonality of species by site and area, we expect that temporal changes in diversity, occurrence, and abundance could be assessed, which would be valuable for tracking the effects of climate change, fishing, management and conservation strategies.

Given the prevalence of divers seeing sharks, and that both common and rare species are preferred by divers (Huveneers et al., 2017) there is cause to consider conservation and protection measures that ensure longterm sustainability of these species for the industry. This could be particularly important given that the sharks reported in the GFSC match those caught by artisanal and subsistence fishers in the region (Glaus et al., 2015). The key to successfully working with fishing groups towards conservation could be that the majority of the shark catch, which was made by subsistence fishers for local consumption, matches the species associated with dive industry. This means that both the dive tourism industry, and subsistence fishers depend, at least in part, on successfully managed and protected shark populations. We suggest there is a need for strong collaborative efforts between the dive tourism industry and subsistence fisheries, community groups, and managers to effectively manage sharks for long-term healthy populations, such as through commitments made by Fiji to meet national and international conservation goals.

Our results may be key for conservation, management, and policy decisions in Fiji. Fiji has a long history of exploitation and many species have been depleted (Glaus et al., 2015). In response, Fiji has undergone significant changes to management, policy, science, and human use patterns to protect many of these species. At the time of writing, for example, the Ministry is collating available data on shark populations and is committed to shark and ocean conservation through the Convention on Biological Diversity as reflected in the National Biodiversity Strategic Action Plan (HS personal observation) and SDGs. Another feature that has been locally important is the use of traditional or locally managed marine areas, which were previously agreed upon by the few active fishers, chiefs, and communities, and which was subsequently weakened with an increase in populations (Dulvy et al., 2004; Glaus et al., 2015). Recently, however, there have been movements towards identifying unique and special biophysical places that are recognized by local communities (Sykes et al., 2018), and strengthening and supporting locally managed marine areas (Jupiter et al., 2014) to meet various goals to protect sharks and biodiversity (Sykes et al., 2018; Wendt et al., 2018). In many areas, with sharks as a proxy for ecosystem health and management needs, our results may provide the best information for the design of protected areas and conservation strategies, and our methodologies may be adaptable for monitoring these management strategies.

### 4.5 Impact

It is widely accepted that public awareness and literacy about the oceans can improve the environment and conservation successes (Costa & Caldeira, 2018; Guest et al., 2015). Involving community leaders and the public in the design of management plans can promote trust and improve legitimacy of management strategies (Dehens & Fanning, 2018; Tonin, 2018). Further, involving the public in data collection, such as through citizen science programs, can improve conservation through local stewardship (Cooper et al., 2007; McKinley et al., 2017). Taken together, building scientific programs that provide opportunities for literacy and participation have added advantages for long-term ocean conservation success.

Our study arose from the need of dive operators to document sharks but reached an international public audience. This multi-year, cross country census of hundreds of dive sites, in collaboration with 39 dive operators, allowed the program to reach divers visiting Fiji from around the world. This reach has value in itself. Although we did not document data about the individual contributors, a previous study estimated that 63,000 divers visit Fiji annually from around the world (Vianna et al., 2011). From these numbers, we estimate that the GFSC captured more than half of all dives made across the country during each census, which suggests that our mission to document these animals may have had a broader education and awareness outcome. It would be ideal to capture this information in future studies, but here the team prioritized simplicity and privacy and opted not to collect personal information.

Programs, like the GFSC, eOceans, and other broad citizen science projects and platforms provide numerous opportunities for individuals and communities to collaborate around mutual interests. In Fiji, approximately 49,000 people (78% of all divers) are engaged in shark diving each year (Vianna et al., 2011) and this provided an opportunity for the dive tourism industry, operators, guides, and scientists to team up to document sharks. Using sharks, we were able to also gather data on other species (e.g., rays and turtles), which are even more data poor in Fiji (not analyzed here). The GFSC also provided an opportunity for cross-community, cross-sector, cross-interest collaboration and has sparked new relationships for future research endeavors.

Finally, each of these additional outcomes, including increasing the number of education and outreach touch point, collaboratively reporting and collecting data, and promoting discussion, awareness, and literacy at the community level and beyond has the potential to contribute to something much larger than the GFSC itself. Through these cross-community and international conversations, we had the opportunity to discuss biodiversity, the need for predators, concerns for sharks and the oceans locally and globally. These added outcomes of the GFSC can tie into much larger goals, beyond describing animal populations. For example, broad participatory science programs like the GFSC and eOceans can help to meet various Aichi Targets (https://www.cbd.int/sp/targets/), such as Targets 1 and 2, which states that “by 2020, at the latest”, “…people are aware of the values of biodiversity” and “… biodiversity values have been integrated into national and local development and poverty reduction strategies and planning processes … are being incorporated into national accounting… and reporting systems”, or SDG 14.A “…increase scientific knowledge, develop research capacity and transfer marine technology …to improve ocean health and to enhance the contribution of marine biodiversity”, or the UN Decade of Ocean Science for Sustainable Development which aims to “develop the scientific research and innovative technologies that can connect ocean science with the needs of society”. In future projects, efforts are needed to begin to measure the success towards these goals.

### 5. Conclusions

Our study provides the first at-sea account of where these eleven shark species are spending at least part of their time, and therefore adds new insights on the distribution and relative abundance. By aggregating the observations of expert and novice divers (i.e., tourist divers assisted by their dive guides) the Great Fiji Shark Count created an extensive longitudinal dataset on hundreds of dive sites across Fiji. Although there are limitations associated with this type of data the extent of sampling demonstrates the value of collaborative citizen science programs for filling data and knowledge gaps on marine megafauna, and the power of a community to come together to assess the issues that matter to them. These results may be used to prompt new scientific questions, to evaluate or resolve management and policy strategies, or as a contemporary baseline that future populations and human use patterns may be compared against. As well, citizen science programs of this magnitude considerably increase the number of opportunities to exchange knowledge, experience, and ideas between industry and science, while also providing frequent touch points with the broader community (e.g., tourists) to promote education and outreach opportunities on issues affecting the oceans. These social outcomes may help meet various sustainable development and biodiversity conservation goals, and deserve further inclusion in scientific and management discussions.

## Acknowledgements

We thank eOceans for data collection, storage, management, analytics, and dissemination, and for in-kind contribution of CWP’s time; Marine Ecology Consulting for facilitating, data entry, and collation the GFSC; and M Neumann for initiating and supporting the GFSC. CWP thanks O MacKenzie for support. The following dive shops and operators contributed organizational support and/or observations to the GFSC: Adrenalin Watersports, Aquatrek Beqa, BAD – Beqa Adventure Divers, BAD – Projects Abroad, Barefoot Kuata, Barefoot Manta, Captain Cook Daycruise, Castaway, Dive Adrenalin, Dive Tropex Tokoriki, Diveaway – Hideaway, Dolphin Bay, Frontier Fiji, GVI – Global Vision International, Koro Sun, L’Aventure Dive Centre, Jean Michel Cousteau Resort, Lalati Resort and Spa, Makaira Resort, Mamanucas Environment Society, Matava Eco Resort-Mad Fish Dive Centre, Nai’a Cruises, Naigani Island, Navini Is Resort, Pacific Harbour Fishing Group, Paradise Taveuni, Ra Divers, Reef Safaris – Barefoot Lodge, Reef Safaris (Bounty Island), Reef Safaris (Intercontinental), Reef Safaris (South Seas Island), Reef Safari – Shangri La Fijian Resort, Reef Safaris (Tivua Island), Safari Lodge, School Marine Sciences University South Pacific, SPAD, Storck Cruises, Taveuni Dive, Taveuni Ocean Sports, Tokoriki Diving, Viti Watersports, Volivoli Beach, Waidroka Bay Resort, Wakaya Club & Spa, Wananavu. We also thank A Batibasaga, N Ledua, S Gow, K Brown and C Cokanisiga for logistical support, and The Great Fiji Shark Count sponsors Beqa Adventure Divers, Fiji Ministry of Fisheries, Marine Ecology Consulting (Fiji), Ocean Soaps (Punjas Fiji), Project Aware, Save Our Seas Foundation, Shark Foundation Switzerland, Shark Savers, WWF South Pacific Programme.

